# ZFP36L1 regulates *Fgf21* mRNA turnover and modulates alcoholic hepatic steatosis and inflammation in mice

**DOI:** 10.1101/2021.05.11.443631

**Authors:** Chandra S. Bathula, Jian Chen, Perry J. Blackshear, Yogesh Saini, Sonika Patial

**Author notes:** **Corresponding author:** *Sonika Patial, DVM, PhD, DACVP*, Department of Comparative Biomedical Sciences, School of Veterinary Medicine, Louisiana State University, Baton Rouge, LA 70803, USA.

## Abstract

Zinc finger protein 36 like 1 (ZFP36L1) enhances the turnover of mRNAs containing AU-rich elements (AREs) in their 3’untranslated regions (3’UTR). The physiological and pathological functions of ZFP36L1 in liver, however, remain largely unknown. To investigate the role of ZFP36L1 in liver physiology and pathology, we generated liver-specific ZFP36L1-deficient (Zfp36l1^flox/flox^ /Cre^+^; L1^LKO^) mice. Under normal conditions, the L1^LKO^ mice and their littermate controls (Zfp36l1^flox/flox^/Cre^-^; L1^FLX^) appeared normal. When fed a Lieber-DeCarli liquid diet containing alcohol, L1^LKO^ mice were significantly protected from developing alcohol-induced hepatic steatosis and inflammation compared to L1^FLX^ mice. Serum ALT levels were significantly increased in alcohol-fed L1^FLX^ versus alcohol-fed L1^LKO^ mice. RNA-Seq analysis revealed 584 differentially-expressed transcripts in L1^FLX^ alcohol-fed mice, many of which were inflammatory mediators, compared to only 159 in alcohol-fed L1^LKO^ mice. Most importantly, fibroblast growth factor 21 (*Fgf21*) mRNA was significantly increased in the livers of alcohol-fed L1^LKO^ mice but not in the alcohol-fed control group. The *Fgf21* mRNA contains three AREs in its 3’UTR, and *Fgf21* 3’UTR was directly regulated by ZFP36L1 in luciferase reporter assays. Steady state levels of *Fgf21* mRNA were significantly decreased by wildtype ZFP36L1, but not by a non-binding zinc-finger ZFP36L1 mutant. Finally, wildtype ZFP36L1, but not the ZFP36L1 mutant, bound to *Fgf21* 3’UTR ARE RNA probe. Our results demonstrate that ZFP36L1 inactivation protects against alcohol-induced hepatic steatosis and liver injury, possibly by stabilizing *Fgf21* mRNA. Our findings suggest that the modulation of ZFP36L1 may be beneficial in the prevention or treatment of human alcoholic liver disease.

## INTRODUCTION

The mouse zinc finger protein 36 (ZFP36 or TTP) family consists of four RNA binding proteins, i.e., zinc finger protein 36, also known as tristetraprolin (TTP), zinc finger protein 36 like 1 (ZFP36L1), zinc finger protein 36 like 2 (ZFP36L2), and zinc finger protein 36 like 3 (ZFP36L3). All four of these proteins regulate mRNA stability by binding to AU-rich elements (AREs) within the 3’-untranslated regions (3’UTRs) of target mRNAs, resulting in ARE-mediated mRNA degradation (1, 2). Despite their common mode of biochemical action, the ZFP36 family members appear to play roles in different physiological processes. This is apparent from the fact that germline deletion of all four ZFP36 family members in mice has resulted in varied phenotypes (3–6). ZFP36 or TTP is the best-studied family member, and plays an essential role in control of inflammation (7, 8). However, the roles of the other three members of the ZFP36 family remain poorly understood.

Recent studies have found that ZFP36L1 is involved in the regulation of mRNAs associated with metabolic functions. For example, ZFP36L1 has been shown to post-transcriptionally regulate the levels of the low-density lipoprotein receptor (LDLR), a protein that is highly expressed in the liver and is a major determinant of circulating levels of LDL, linking ZFP36L1 to cholesterol metabolism (9). ZFP36L1 has also been shown to post-transcriptionally regulate uncoupling protein 1 (UCP1), which plays an important role in thermogenesis (10). Along similar lines, ZFP36L1 was recently shown to regulate liver bile acid metabolism (11). These findings indicate that ZFP36L1 may play critical roles in regulating liver metabolic homeostasis. However, the detailed physiological functions of ZFP36L1 remain particularly elusive, partly due to the embryonic lethal phenotype that is observed following germline deletion of ZFP36L1 in mice (4).

In this study, we sought to understand the physiological roles of ZFP36L1 in the liver. We hypothesized that the removal of ZFP36L1 could alter certain pathological processes associated with liver. To test this, we employed Cre-lox technology to delete *Zfp36l1* in hepatocytes and cholangiocytes (12). We then subjected liver-specific ZFP36L1-deficient (L1^LKO^) and their littermate control mice (L1^FLX^) to a model of alcoholic steatohepatitis (13). The mice were analyzed for histological, biochemical, and transcriptional alterations associated with the development of alcoholic steatosis, liver injury, and inflammation. The results show a novel protective effect of liver-specific ablation of ZFP36L1 against alcohol-induced steatosis and liver inflammation, and identified *Fgf21* mRNA as a likely target of ZFP36L1.

## MATERIALS AND METHODS

### Mice

The floxed mice carrying loxP-flanked exon 2 of *Zfp36l1* alleles were generated by gene targeting in C57BL/6 embryonic stem cells at Xenogen Biosciences (Cranbury, NJ). Liver-specific *Zfp36l1* knockout (*Zfp36l1*^flox/flox^/Cre^+^; L1^LKO^) mice were generated by crossing the homozygous floxed mice (*Zfp36l*^flox/flox^) with albumin-Cre (Alb-Cre) (14) mice (Jackson Laboratory, Bar Harbor, Maine). Genotype status of the progeny was determined by polymerase chain reaction (PCR) using primer 1 (P1), 5’-CTCTTGCTGGTCACTACCGTCGCT-3’ and primer 2 (P2), 5’-TCAATGTAAGCCAGCAAGTGCAGC-3’, which amplified a 484 bp endogenous *Zfp36l1* WT allele and a 618 bp mutant *Zfp36l1* floxed allele. PCR conditions were: 93°C for 3 min followed by 30 cycles of 93°C for 30 s, 58°C for 30 s, and 72°C for 1 min, with a final extension of 72°C for 10 min. Eight to twelve-week-old female L1^LKO^ and littermate control (Zfp36l1^flox/flox^/Cre^-^; L1^FLX^) mice were used in all studies. All the animal experiments were performed in accordance with principles and procedures outlined in the National Institute of Health Guide for the Care and Use of Laboratory Animals and were approved by the Louisiana State University Animal Care and Use Committee.

### Lieber-DeCarli liquid diet model of chronic-plus-binge model of alcoholic liver disease

8-12 week old female L1^LKO^ or L1^FLX^ mice were subjected to alcoholic liver disease as described by Bertola et al (13). Briefly, mice were first acclimatized for 5 days to tube feeding and liquid diet *ad libitum*. Following acclimatization, mice were maintained for 10 days on a fresh liquid diet (free access) containing 5% ethanol (vol/vol) (Lieber-DeCarli ‘82 Shake and Pour ethanol liquid diet; product number F1258SP; Bio-Serv) or pair-fed with a control isocaloric diet (Lieber-DeCarli ‘82 Shake and Pour control liquid diet; product number F1259SP; Bio-Serv) in which the same number of calories was obtained from dextrin maltose. The freshly constituted liquid diet was stored in the refrigerator and used within three days. The refrigerated diet was brought to room temperature, and a calculated volume of fresh ethanol was added daily. On day 11, a single intragastric (oral gavage) administration of ethanol (5 g ethanol/Kg body weight; 31.5% ethanol) or an isocaloric dose of maltose-dextrin (control group) was administered between 8 AM and 9 AM. Mice were humanely euthanized 9 hours after gavage feeding, and serum and liver were collected.

### Necropsy and tissue collection

Mice were humanely euthanized, midline laparotomy was performed, and blood was collected by cardiac puncture for serum. The livers were removed and weighed, and the left lateral lobe was dissected into three parts, of which the first part was stored in RNA later (Life Technologies, Carlsbad, CA), the second part was snap-frozen and stored at −80°C, and the third part was fixed in 10% neutral buffered formalin (NBF). The remaining liver lobes were fixed in 10% NBF.

### Histology

5 μm sections of liver were stained with Hematoxylin and Eosin (H&E) for routine histological analyses. Quantification of steatosis was performed by stereological point counting (SPC) (15). Briefly, the histological microscopic images were overlaid by a regular grid of points. The grid contained approximately 300 intersections within the tissue boundary that were counted as either fat deposits or no fat deposits depending on whether they covered the fat droplets or not. The fractional area of fat droplets was determined by dividing the number of points covering fat droplets to the total number of points.

### Serum enzymes and lipids analyses

Analyses for ALT, AST, cholesterol, and triglycerides were performed on a Beckman Coulter AU680 chemistry analyzer using Beckman Coulter reagents (Beckman-Coulter, Irving, TX) at the clinical pathology core at LSU.

### ELISA

Serum levels of FGF21 were determined using an ELISA kit (R&D systems, Minneapolis, MN) according to the manufacturer’s instructions. The Optical Density (OD) was measured at 450 nm and 570 nm using Spark® Multimode Microplate Reader (Tecan Trading AG, Mannedorf, Switzerland). The OD readings at 570 nm were subtracted from those at 450 nm to correct for optical imperfections in the plate.

### RNA extraction

Total RNA was extracted using the GE Healthcare Illustra RNAspin MiniRNA isolation kit, according to the manufacturer’s instructions (GE Healthcare, Little Chalfont, UK). The RNA content and purity were determined by measuring absorbance at 260 and 260/280 nm, respectively, on a Nanodrop™ 8000 spectrophotometer (Thermo Fisher Scientific, Waltham, MA). The quality of the RNA was determined using an Agilent Bioanalyzer 2100 (Agilent, Santa Clara, CA).

### Quantitative RT-PCR

Real-time PCR for the quantification of *Tnf, Mcp1, Il10, and Cd36* transcripts was performed using the SYBR Green method. Briefly, 1μg of total RNA was used for cDNA synthesis using iScript cDNA synthesis kit (Bio-Rad, Hercules, CA). Quantitative real-time RT-PCR was performed on the ABI Prism 7900 Sequence Detection System (Applied Biosystems). Primer sequences were: *Tnf*, **Forward** 5’-CTTCTGTCTACTGAACTTCGGG-3’, **Reverse** 5’-CAGGCTTGTCACTCGAATTTTG-3; *Mcp1*, **Forward** 5’-GTCCCTGTCATGCTTCTGG-3’, **Reverse** 5’-GCTCTCCAGCCTACTCATTG-3’; *Il10*, **Forward** 5’-AGCCGGGAAGACAATAACTG-3’, **Reverse** 5’-GGAGTCGGTTAGCAGTATGTTG-3’; *Cd36*, **Forward** 5’-GCGACATGATTAATGGCACAG-3’, **Reverse** 5’-GATCCGAACACAGCGTAGATAG-3’. The ct values were normalized to *Gapdh* internal controls. The expression levels were calculated according to the 2^−ΔΔct^ method. Real-time PCR for *Fgf21* and *Zfp36l1* was performed using Taqman assays (*Fgf21*:hs00173927; *Zfp36l1*: hs00245183).

### Library preparation and RNA deep sequencing

Library preparation and RNA sequencing was performed at Novogene (www.en.novogene.com). Briefly, 1 μg of RNA was used for cDNA library construction using an NEBNext® Ultra 2 RNA Library Prep Kit for Illumina® (New England Biolabs, Ipswich, MA) according to the manufacturer’s protocol. mRNA was enriched using oligo (dT) beads followed by two rounds of purification and random fragmentation. The first strand cDNA was synthesized using random hexamers followed by generation of the second-strand. After a series of terminal repair, poly-adenylation, and sequencing adaptor ligation, the double-stranded cDNA library was completed following size selection and PCR enrichment. The resulting 250-350 bp insert libraries were quantified using a Qubit 2.0 fluorometer (Thermo Fisher Scientific, Waltham, MA) and quantitative PCR. Size distribution was analyzed using an Agilent 2100 Bioanalyzer (Agilent Technologies, Santa Clara, CA). Qualified libraries were sequenced on an Illumina Nova Seq 6000 Platform (Illumina, Inc., San Diego, CA) using a paired-end 150 run (2×150 bases). Approximately 20 M raw reads were generated from each library.

### Immunoblotting

Cells/tissues were lysed using Pierce™ RIPA buffer (Thermo Fisher Scientific, Waltham, MA) supplemented with Pierce™ protease inhibitor mixture (Thermo Fisher Scientific, Waltham, MA) and phosphatase inhibitors (10 mM sodium fluoride and 1 mM sodium orthovanadate). Tissues were mechanically homogenized using a homogenizer (Fisherbrand™ 150 handheld Homogenizer, Thermo Fisher Scientific, Waltham, MA). Tissue lysates were centrifuged (13,000 x g, 10 min, 4°C) to remove insoluble material, and protein concentration of the supernatants were measured using the Bradford assay (Bio-Rad Laboratories, Hercules, CA). Denatured proteins were separated on a 4-12% Bis-Tris plus gel (Invitrogen, Carlsbad, CA), transferred onto PVDF membranes (Invitrogen, Carlsbad, CA), and probed with a 1:1000 dilution of BRF1/2 primary antibody (Cell signaling Technology, Danvers, MA), followed by a 1:3000 dilution of HRP-conjugated goat anti rabbit IgG. Signals were determined either on a ChemiDoc Imaging System (Bio-Rad Laboratories, Hercules, CA) or on X-ray film.

### Luciferase reporter assay

HEK293 cells were procured from ATCC. Cells were cultured in EMEM with 10% FBS. pcDNA 3.1+/C-(K)-DYK *Zfp36l1*, pcDNA 3.1+/C-(K)-DYK *Zfp36l1*-zinc finger mutant C135R, C173R (two amino-acid mutations within the two zinc fingers at position 135 and 173), and pcDNA 3.1+/C-(K)-DYK empty vector were obtained from GenScript Biotech (GenScript, Piscataway, NJ). Ambion^R^ pMIR-REPORT™ Luciferase expression reporter vector was obtained from Thermo Fisher Scientific (Thermo Fisher Scientific, Waltham, MA), and the 3’UTR of *Ffg21* mRNA was cloned into pMIR-REPORT Luciferase reporter vector. Transfections were done with Lipofectamine reagent (Invitrogen Carlsbad, CA). Luminescence was determined using Promega Luciferase Reporter Assay Kit (Promega, Madison, WI), according to the manufacturer’s instructions, on a Spark® Multimode Microplate Reader (Tecan Trading AG, Mannedorf, Switzerland). Briefly, cells were seeded onto 6 well plates and transfected with the respective constructs (0.5 μg) when they reached a confluency of approximately 60-65%. 48 hours post-transfection, growth medium was removed, and cells were gently rinsed with PBS, followed by addition of 1X reporter lysis buffer (RLB) to lyse the cells. To ensure complete lysis, cells were subjected to a single freeze-thaw cycle. Lysed cells were scraped, transferred into microcentrifuge tubes, and centrifuged at 12,000 x g for 15 seconds at room temperature. The supernatants were transferred into new microcentrifuge tubes. 20 μL of cell lysate was used per well of a 96 well plate for luciferase assays. The Spark® Multimode Microplate Reader was programmed to add 100 μL of luciferase assay reagent per well with the injector. Luminescence was determined immediately (10 sec). Luminescence was normalized to Beta-galactosidase measurements.

### Co-transfection assay

For co-transfection assays, HEK293 cells were co-transfected with pcDNA 3.1+/C-(K)-DYK empty vector alone, the pcDNA 3.1+/C-(K)-DYK *Zfp36l1*-WT plasmid or the zinc-finger mutant, pcDNA 3.1+/C-(K)-DYK *Zfp36l1*-C135R, C173R, and the full length pcDNA 3.1+/C-(K)-DYK-*Fgf21* plasmid, which contained an intact 3’UTR (GenScript, Piscataway, NJ). Two concentrations of *Zfp36l1* (WT and mutant) plasmids were used, 0.25 μg and 0.50 μg, while 0.5 μg of the *Fgf21* plasmid was transfected in all. The amount of total transfected plasmid was kept constant for all the plates, and normalized by the addition of vector control plasmid. The cells were harvested after 48 hours, RNA was isolated, cDNA was prepared using iScript™ Reverse Transcription Supermix (Bio-rad, Hercules, CA), and the levels of *Fgf21* and *Zfp36l1* mRNAs were assessed by quantitative real-time PCR.

### Electrophoretic Mobility Shift Assays (EMSA)

RNA-ARE probes were designed from the *Fgf21*-3’UTR ARE sequence and synthesized with 5’-biotin end-labelling (IDT DNA technologies). HEK293 cells were transfected with the ZFP36L1 expression plasmid (pcDNA 3.1+/C-(K)-DYK *Zfp36l1*) or its zinc finger mutant (pcDNA 3.1+/C-(K)-DYK *Zfp36l1*-C135R, C173R), and cytosolic cell extracts were prepared. Briefly, the transfection mixture was removed and cells were rinsed with ice cold Ca^2+^ and Mg^2+^ free DPBS. The cells were then scraped and centrifuged at 600 x g for 3 min at 4°C. The supernatants were removed, and the pellets were rinsed with ice cold DEPC-treated water containing 8 μg/ml leupeptin, 0.2 mM phenylmethylsulfonyl fluoride, and 1 μg/ml pepstatin A. The cells were then lysed in a hypotonic cell lysis buffer containing 10 mM HEPES (pH 7.6), 5 mM KCl, 5% (v/v) glycerol, 0.25% (v/v) Nonidet P-40, 1 μg/ml pepstatin A, 0.1 mM phenylmethylsulfonyl fluoride, and 8 μg/ml leupeptin. Lysates were centrifuged for 15 min at 15,000 × *g* at 4 °C. The supernatant was transferred to a fresh tube, and KCl and glycerol were added to final concentrations of 40 mM and 15% (v/v), respectively. Cell extracts were aliquoted and stored at −80 °C until ready to use. For the EMSA assay, approximately 8 μg of cytosolic extracts prepared from HEK293 cells transfected with either the vector alone, or with the wt or mutant ZFP36L1 expression constructs, were incubated with 300 nM 5’-biotin-end labelled ARE probes at room temperature for 30 min. The total reaction volume was 20 μl and consisted of 10 mM HEPES (pH 7.6), 40 mM KCl, 2.5% (v/v) glycerol, 3 mM MgCl2, 2.5 μg/μl heparin, and 50 ng/μl yeast tRNA. Following incubation, the reaction mixture was loaded onto 6-8% non-denaturing acrylamide gels and subjected to electrophoresis (120V for 60 min in 0.5 X TBE buffer). Gels were transferred onto Biodyne B nylon membranes (0.45 μm; Thermo Fisher Scientific, Rockford, IL, USA) in 0.5X TBE buffer (60V for 30 min). RNA-protein complexes and the unbound RNA probe were detected using the Chemiluminescent Nucleic Acid Detection Module (Thermo Scientific), following manufacturer’s instructions.

### Statistical Analysis

Statistical significance between two groups was determined by Student’s *t* test assuming unequal variance. Significant differences among groups were determined by one-way analysis of variance (ANOVA) followed by Tukey’s post hoc test for multiple comparisons. All data were expressed as mean ± SEM. A *p* value <0.05 was considered statistically significant. Statistical analyses were performed using GraphPad Prism 7.0 (GraphPad Software, La Jolla, CA).

### Nucleotide sequence accession number

The RNA-Seq data have been deposited in the NCBI GEO under accession number **GSE163444**

## RESULTS

### Generation and characterization of ZFP36L1 liver-specific knockout (LKO) mice

To investigate the role of ZFP36L1 in liver, liver-specific *Zfp36l1*-deficient mice (L1^LKO^) were generated using the Cre-LoxP recombination approach. RNA-Seq analysis of liver RNA showed at least a 5-fold reduction in the mRNA levels of *Zfp36l1* in L1^LKO^ mice compared to littermate control L1^FLX^ mice (**Figure 1A**). Western blot analysis of liver tissue lysates showed minimal to no detectable expression of ZFP36L1 in L1^LKO^ mice compared to littermate control L1^FLX^ mice (**Figure 1B**). Quantitatively, the expression of ZFP36L1 was significantly reduced in L1^LKO^ mice compared to littermate control L1^FLX^ mice (**Figure 1C**). The L1^LKO^ mice developed normally and did not exhibit any growth defects or anatomical or histological abnormalities (data not shown).

**Figure 1:**
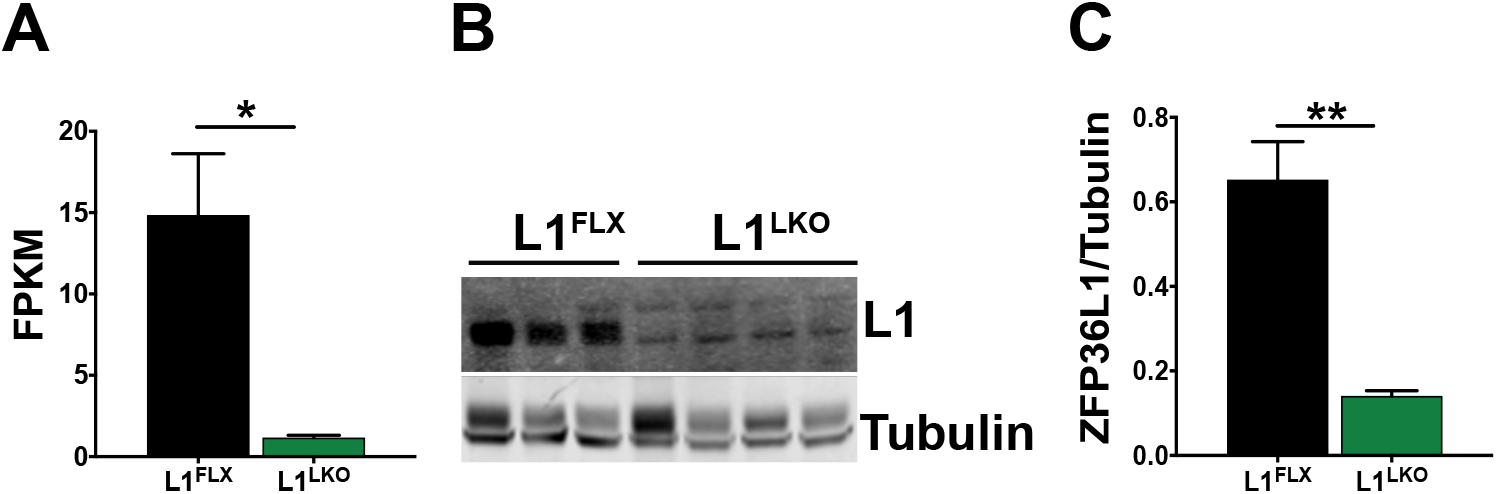
mRNA and protein expression of ZFP36L1 in control (L1^FLX^) versus ZFP36L1 liver-specific knockout (L1^LKO^) mice. **A.** FPKM (Fragments Per Kilobase of transcripts per Million mapped reads) values from liver RNA-Seq data showing the expression of *Zfp36l1* mRNA in liver homogenates from ZFP36L1-sufficient (L1^FLX^) and liver-specific ZFP36L1-deficient (L1^LKO^) mice. n=4. Data are represented as mean ± SEM). **B.** An immunoblot showing expression of ZFP36L1 in liver homogenates from ZFP36L1-sufficient (L1^FLX^) and liver-specific ZFP36L1-deficient (L1^LKO^) mice. n=3 for L1^FLX^; n=4 for L1^LKO^ mice. **C.** Image J quantification of band intensity in immunoblot shown in **B**.

### Alcohol-fed L1^LKO^ mice exhibit significantly attenuated hepatic steatosis, injury, and inflammation compared to alcohol-fed littermate control L1^FLX^ mice

In order to determine the effect of ZFP36L1 deletion in stressed liver cells, we subjected L1^FLX^ and L1^LKO^ mice to alcohol-induced liver injury and steatosis by feeding them the alcohol-containing Lieber-DeCarli diet (alcohol-group). The weights of both the L1^FLX^ and L1^LKO^ mice did not change significantly post alcohol ingestion over a period of 11 days compared to their initial body weights (**Supplemental Figure 1A**). Alcohol significantly increased the liver weight to body weight ratio after alcohol-diet feeding in both L1^FLX^ and L1^LKO^ mice, but the difference was not significant between the alcohol-fed L1^FLX^ and L1^LKO^ groups (**Figure 2A**).

**Figure 2:**
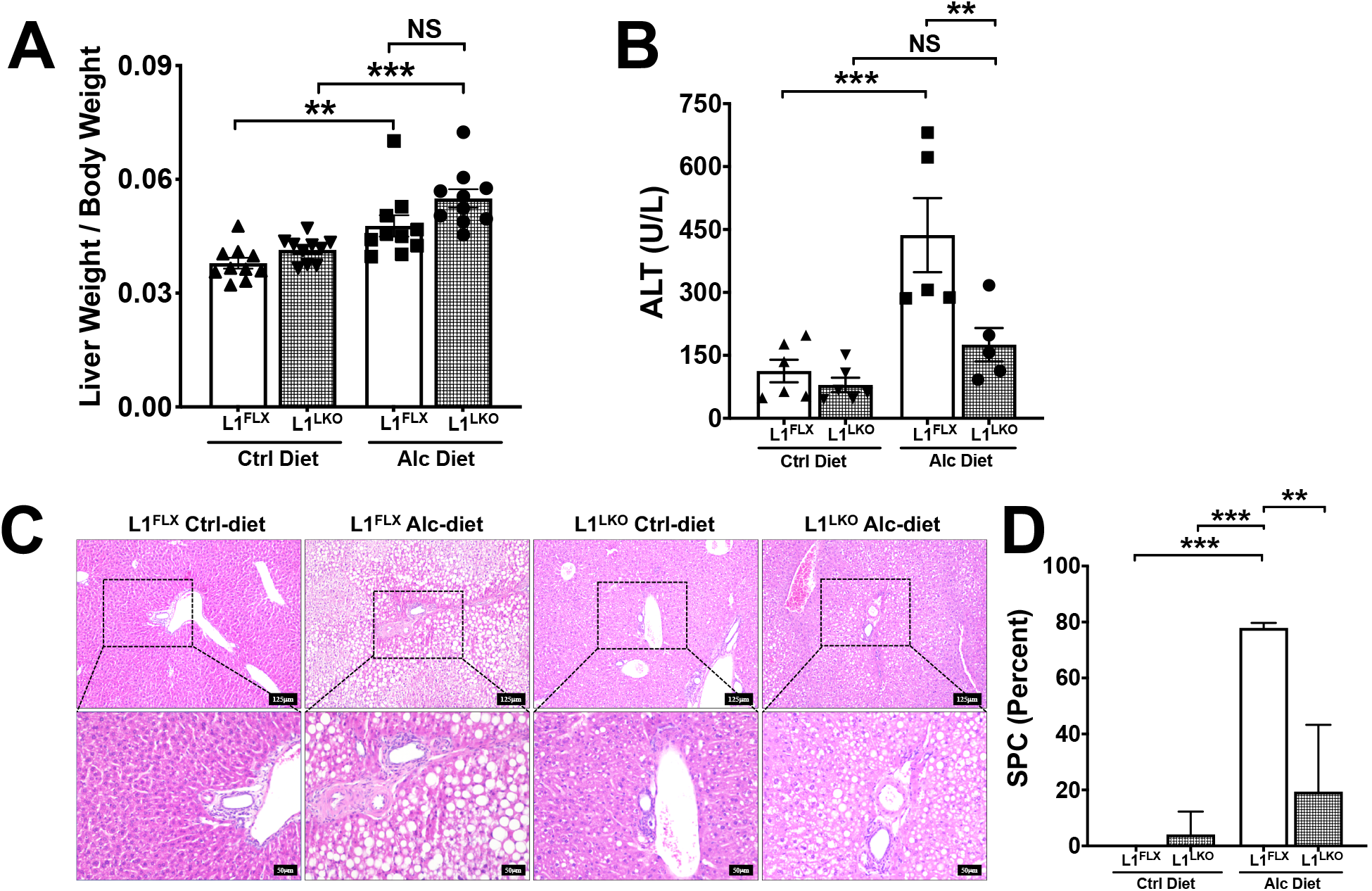
ZFP36L1 LKO mice were protected from alcohol-induced liver injury. **A.** Liver weight to body weight ratios of alcohol-diet-fed ZFP36L1-sufficient (L1^FLX^) and liver-specific ZFP36L1-deficient (L1^LKO^) mice and those that were pair-fed with control diet. n=10 in each group. **B.** Serum levels of ALT in alcohol-diet-fed ZFP36L1-sufficient (L1^FLX^) and liver-specific ZFP36L1-deficient (L1^LKO^) mice and the control-diet-fed ZFP36L1-sufficient (L1^FLX^) and liver-specific ZFP36L1-deficient (L1^LKO^). n=5 (alcohol-diet group); n=6 (control-diet group). Statistical analysis was performed by one-way ANOVA followed by Tukey’s correction for multiple comparisons. All data are shown as mean ± SEM. **p<0.01; ***p<0.001. **C.** Representative liver histology images from ZFP36L1-sufficient (L1^FLX^) and liver-specific ZFP36L1-deficient (L1^LKO^) mice that were fed with alcohol-diet versus those that were pair-fed with control-diet are shown. **D.** Quantification of hepatic steatosis of images shown in **C**.

Alcohol results in hepatic lesions, particularly steatosis, hepatic injury, and hepatic inflammation (16). Serum levels of alanine aminotransferase (ALT), a marker of liver injury, were significantly increased in alcohol-fed L1^FLX^ mice compared to control-diet-fed (pair-fed) L1^FLX^ mice. However, alcohol ingestion did not increase the levels of ALT in L1^LKO^ mice (**Figure 2B**). The levels of aspartate aminotransferase (AST) were also increased in alcohol-fed mice but the difference was not statistically significant between L1^FLX^ and L1^LKO^ groups (**Supplemental Figure 1B**). The serum levels of triglycerides and cholesterol were not significantly increased post alcohol-diet feeding compared to control-diet feeding in either of the two genotype groups (**Supplemental Figure 1C-D**).

Steatosis is the earliest response of the liver to alcohol ingestion and is characterized by the accumulation of lipid droplets within the hepatocytes, starting at the centrilobular areas and progressing towards the mid-lobular and peri-portal regions in later stages (16, 17). Consistent with these reports, the alcohol-diet significantly increased microvesicular and macrovesicular steatosis in L1^FLX^ mice compared to control-diet fed L1^FLX^ mice. Remarkably, alcohol-diet-fed L1^LKO^ mice exhibited markedly diminished hepatic steatosis compared to the alcohol-diet-fed L1^FLX^ mice (**Figure 2C-D**).

Alcohol-induced steatosis is usually accompanied by hepatic inflammation (16). Certain pro-inflammatory mediators, particularly TNF and MCP1, have been implicated in alcoholic liver inflammation (18, 19). Similarly, IL-10 and CD36, a fatty acid translocase, has also been shown to play a role in alcoholic liver disease (20),(21). Therefore, we next assessed the mRNA expression levels for these pro-inflammatory cytokines and mediators in alcohol-diet fed L1^FLX^ and L1^LKO^ mice. The mRNA expression levels of both *Tnf* and *Mcp1* were significantly increased in alcohol-diet-fed L1^FLX^ mice compared to control-diet-fed L1^FLX^ mice (**Figure 3A, B**). However, the increase in expression of *Tnf* and *Mcp1* was significantly attenuated in alcohol-fed L1^LKO^ mice compared to alcohol-fed L1^FLX^ mice (**Figure 3A, B**). The mRNA expression levels of *Il10* and *Cd36* also showed similar trends, but the differences were not statistically significant between alcohol-diet-fed L1^LKO^ and alcohol-diet-fed L1^FLX^ mice (**Figure 3C, D**). Taken together, these data demonstrate that liver-specific deletion of ZFP36L1 confers significant protection against hepatic steatosis, hepatic injury, and hepatic inflammation in alcohol-diet-fed mice.

**Figure 3:**
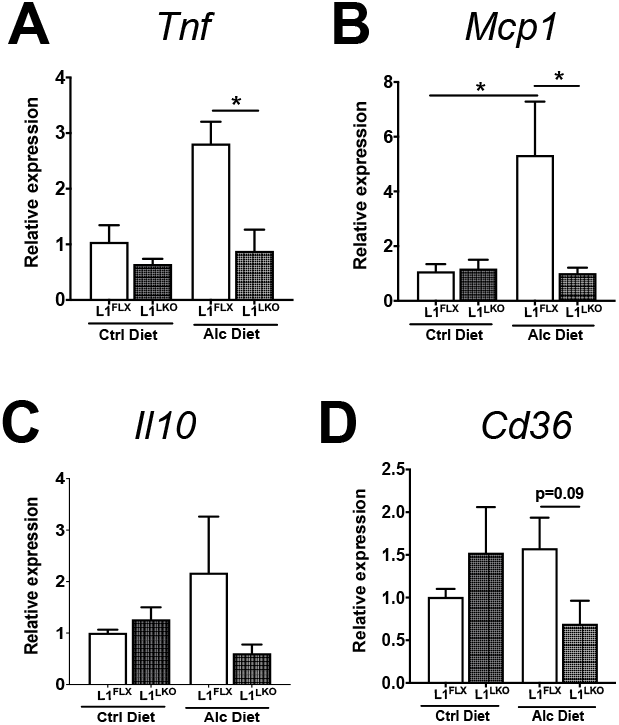
ZFP36L1 LKO mice were protected from alcohol-induced liver inflammation. Relative quantification of *Tnf* (**A**), *Mcp1* (**B**), *Il10* (**C**), and *Cd36* (**D**) mRNA expression in the liver homogenates from alcohol-diet-fed ZFP36L1-sufficient (L1^FLX^) and liver-specific ZFP36L1-deficient (L1^LKO^) mice and those that were pair-fed with control diet. Statistical analyses were performed by one-way ANOVA followed by Tukey’s correction for multiple comparisons. All data are shown as mean ± SEM. n=3-4. *p<0.05, *Tnf*, Tumor necrosis factor; *Mcp1*, Monocyte Chemoattractant Protein-1; *Il10*, Interleukin-10; *Cd36*, Cluster of differentiation 36.

### ZFP36L1 deficiency results in reduced numbers of differentially expressed transcripts in alcohol-fed mice

In order to identify transcriptomic changes through which ZFP36L1 modulates alcoholic hepatic steatosis, hepatic injury, and hepatic inflammation, we performed RNA-Seq analysis of total liver RNA obtained from the four groups: control-diet-fed and alcohol-diet-fed L1^FLX^ and L1^LKO^ mice. The RNA-Seq data are expressed as FPKM (fragments per kilobase per million mapped reads). The inclusion criteria for increase or decrease was set as greater or less than 1.5-fold (log2FC), with adjusted p-values of less than 0.05. Interestingly, while 584 transcripts were significantly differentially expressed (393 up-regulated; 191 down-regulated) in the alcohol-diet-fed L1^FLX^ mice compared to control-diet-fed L1^FLX^ mice, only 159 transcripts were differentially expressed (92 up-regulated; 67 down-regulated) in the alcohol-diet-fed L1^LKO^ mice compared to control-diet-fed L1^LKO^ mice (**Figure 4A and 4B**). Fifty transcripts were differentially expressed (26 up-regulated; 24 down-regulated) between alcohol-diet-fed L1^LKO^ and alcohol-diet-fed L1^FLX^ mice (**Figure 4C**), while only 10 transcripts were differentially expressed (7 up-regulated; 3 down-regulated) between control-diet-fed L1^LKO^ and control-diet-fed L1^FLX^ mice (**Figure 4D**). **Figure 4E** shows a heat map comparing differentially expressed transcripts in the four groups. These data indicate that, in line with the increased steatosis, liver injury, and inflammation seen in the alcohol-diet-fed L1^FLX^ mice, there were larger numbers of transcriptomic changes, whereas minimal to no steatosis and significantly attenuated liver injury and inflammation was associated with comparatively lesser numbers of transcriptomic changes in alcohol-diet-fed L1^LKO^ mice.

**Figure 4:**
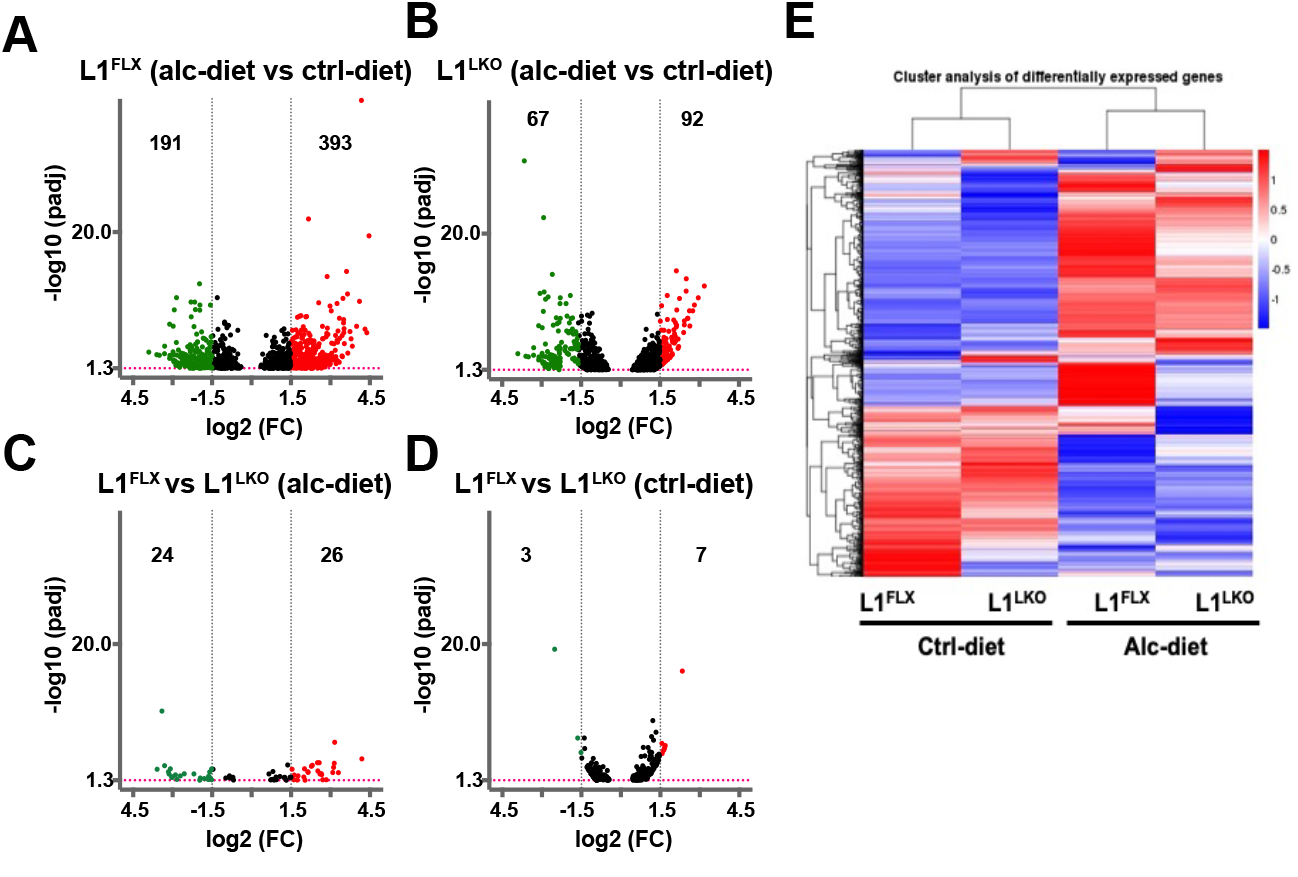
Differentially expressed transcripts in L1^LKO^ mice and L1^FLX^ mice following control-diet and alcohol-diet feeding. Volcano plots depict differentially expressed transcripts (red: upregulated; green: downregulated) in alcohol-diet-fed versus control-diet-fed L1^FLX^ mice (**A**); alcohol-diet-fed versus control-diet-fed L1^LKO^ mice (**B**); alcohol-diet fed L1^FLX^ versus L1^LKO^ mice (**C**); and control-diet-fed L1^FLX^ versus L1^LKO^ mice (**D**). Black vertical dashed lines indicate a log2fold-change cutoff of 1.5. The red horizontal dashed line indicates −log10 (p-value) cutoff of 1.3 (p-value=0.05). Heatmap of differentially expressed transcripts from alcohol-diet-fed ZFP36L1-sufficient (L1^FLX^) and liver-specific ZFP36L1-deficient (L1^LKO^) mice and control-diet-fed ZFP36L1-sufficient (L1^FLX^) and liver-specific ZFP36L1-deficient (L1^LKO^) mice (**E**).

Next, to investigate whether the large numbers of gene expression changes in the alcohol-diet-fed L1^FLX^ versus alcohol-diet-fed L1^LKO^ group could be accounted for by pro-inflammatory mRNAs, we compared the expression levels of known pro-inflammatory mRNAs between the two groups. We found that a large proportion of the known inflammatory genes were significantly up-regulated in the L1^FLX^ group (alcohol-diet-fed L1^FLX^ mice versus control-diet-fed L1^FLX^ mice) but not in the L1^LKO^ group (alcohol-diet-fed L1^LKO^ mice versus control-diet-fed L1^LKO^ mice) (**Table 1**). Key inflammatory genes included toll like 4-receptor *Cd14*, macrophage scavenger receptor, *Marco*, interleukin receptor, *Il3ra*, chemokines including *Ccl2, Ccr1, Cxcl2, Cxcl14*, and *Ccrl2*, TNF receptor superfamily proteins, *Tnfrsf10b* and *Tnfrsf21*, NFKB family proteins *Ikbip* and *Nfkbib*, MAP kinase 4 (*Mapk4*), complement component, *C3ar1*, cell surface receptors, *Cd34, Cd68*, and *Cd44*, protein kinase substrate protein, *Marcksl1*, and metalloelastases, *Mmp12* and *Timp*. In fact, few of the mRNAs that achieved statistical significance in the alcohol-diet-fed L1^LKO^ group were upregulated by a comparatively lower-fold change when compared to the alcohol-diet-fed L1^FLX^ group. These data indicate that alcohol-diet-feeding resulted in significantly increased liver inflammation in L1^FLX^ mice but not in L1^LKO^ mice.

**Table 1:**
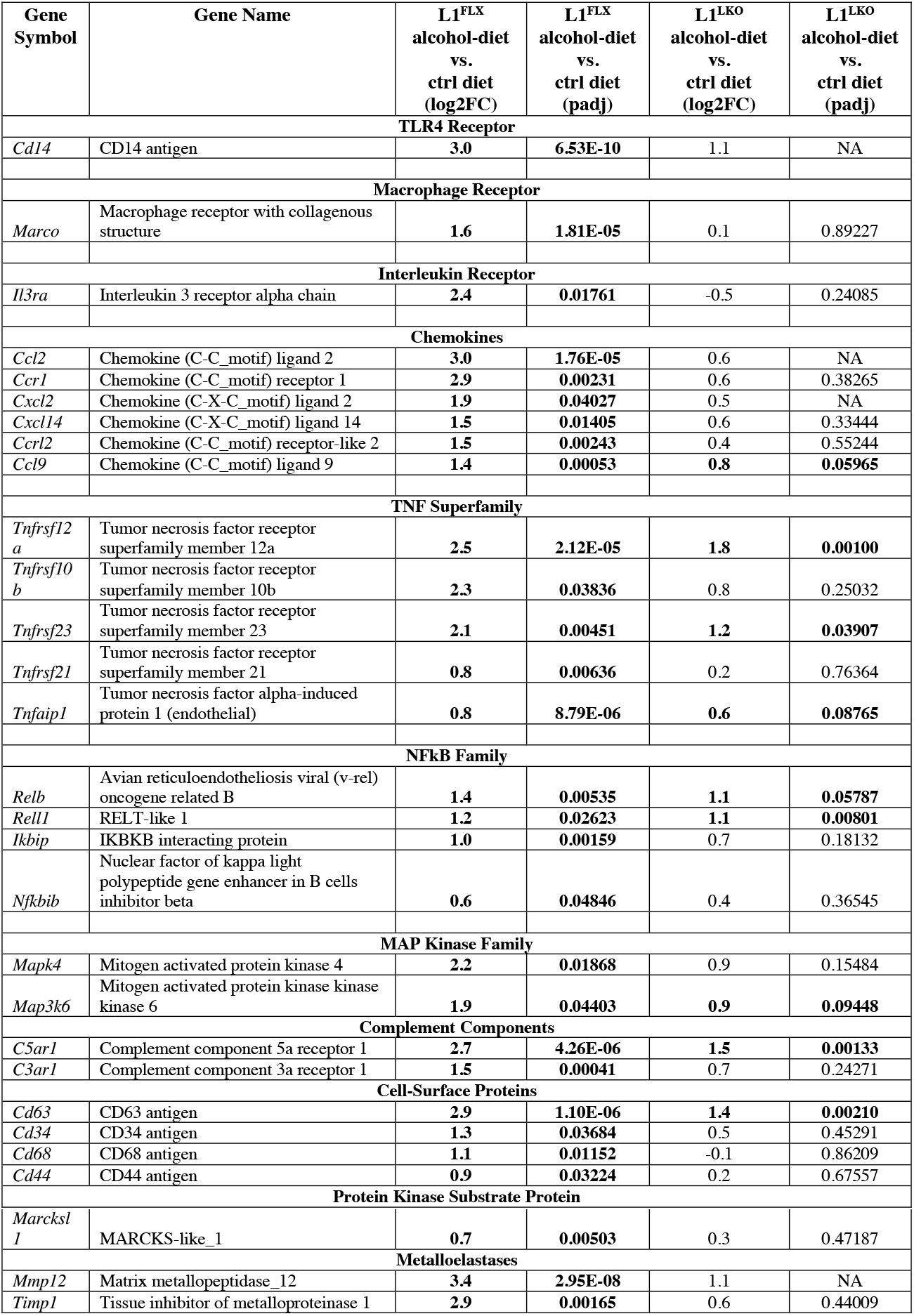
Differentially expressed **inflammatory genes** in alcohol-diet-fed ZFP36L1-sufficient (L1^FLX^) and liver-specific ZFP36L1-deficient (L1^LKO^) mice and control-diet-fed ZFP36L1-sufficient (L1^FLX^) and liver-specific ZFP36L1-deficient (L1^LKO^) mice.

### Comparison of expression of alcohol-regulated metabolic mRNAs in alcohol-diet-fed L1^LKO^ versus the alcohol-diet-fed L1^FLX^ mice

Alcohol ingestion results in metabolic changes including increased lipogenesis, decreased fatty acid oxidation, mobilization of fat from various body deposits into the liver, and impaired VLDL secretion, all resulting in the development of hepatic steatosis (16). A large number of metabolic genes is involved in this process. To delineate the mechanisms for differential responses of the L1^FLX^ versus the L1^LKO^ mice to alcohol-feeding, we queried the RNA-Seq data set for changes in known metabolic transcripts. Sirtuin 1 (*Sirt1*), fatty acid synthase (*Fasn*), stearoyl-coenzyme A desaturase 1 (*Scd*1), Acyl-Coenzyme A dehydrogenase Family Member 8 (*Acad8*), and Cytochrome P450 family 2 subfamily e polypeptide 1 (*Cyp2e1*) were significantly downregulated in alcohol-diet-fed L1^FLX^ group but were unchanged in the alcohol-diet-fed L1^LKO^ group, while Lipin-3 and lipoprotein lipase (*Lpl*) were significantly upregulated in alcohol-diet-fed L1^FLX^ group but unchanged in the alcohol-diet-fed L1^LKO^ group. Peroxisome proliferator activated receptor alpha (*Ppara*) and sterol regulatory element binding transcription factor 1 (*Srebf1*) were significantly downregulated in both the L1^FLX^ and L1^LKO^ groups following alcohol-ingestion, although the fold change was higher in the L1^FLX^ group (**Table 2**).

**Table 2:** Differentially expressed **metabolic genes in** alcohol-diet-fed ZFP36L1-sufficient (L1^FLX^) and liver-specific ZFP36L1-deficient (L1^LKO^) mice and control-diet-fed ZFP36L1-sufficient (L1^FLX^) and liver-specific ZFP36L1-deficient (L1^LKO^) mice.

| Gene Symbol | Gene Name | L1<sup>FLX</sup><br>alcohol-diet<br>vs.<br>ctrl-diet<br>(log2FC) | L1<sup>FLX</sup><br>alcohol-diet<br>vs.<br>ctrl-diet<br>(padj) | L1<sup>LKO</sup><br>alcohol-diet<br>vs.<br>ctrl-diet<br>(log2FC) | L1<sup>LKO</sup><br>alcohol-diet<br>vs.<br>ctrl-diet<br>(padj) |
| --- | --- | --- | --- | --- | --- |
| *Metabolic Regulator genes* |  |  |  |  |  |
| *Sirt1* | Sirtuin 1 | **-0.93** | **0.01** | -0.08 | 0.88 |
| *Sirt3* | Sirtuin 3 | **-1.32** | **0.02** | **-0.90** | **0.001** |
| *Lipid metabolism* |  |  |  |  |  |
| *Lpin1* | Lipin 1 | **-1.12** | **0.07** | **-1.28** | **0.01** |
| *Lpin2* | Lipin 2 | -0.31 | 0.75 | **-0.90** | **0.008** |
| *Lpin3* | Lipin 3 | **1.33** | **0.02** | 0.25 | 0.67 |
| *Lpl* | Lipoprotein lipase | **2.3** | **0.001** | 0.3 | 0.55 |
| *Lipogenesis* |  |  |  |  |  |
| *Srebf1* | Sterol regulatory element binding transcription factor 1 | **-2.28** | **5.99E-05** | **-1.68** | **6.15E-07** |
| *Fasn* | Fatty acid synthase | **-1.78** | **0.05** | -0.64 | 0.32 |
| *Scd1* | Stearoyl Coenzyme A desaturase 1 | **-1.89** | **0.03** | -0.54 | 0.48 |
| *Fatty acid oxidation* |  |  |  |  |  |
| *Ppara* | Peroxisome proliferator activated receptor alpha | **-1.19** | **0.01** | **-0.98** | **0.04** |
| *Acad8* | Acyl-Coenzyme A dehydrogenase Family Member 8 | **-0.65** | **0.03** | -0.29 | 0.36 |
| *Cpt1a* | Carnitine palmitoyltransferase 1a liver | -0.39 | 0.71 | **-0.94** | **0.01** |
| *Cd36* | CD36 antigen | -0.02 | 0.98 | 0.27 | 0.75 |
| *Fgf21* | Fibroblast growth factor 21 | 0.7 | 0.60 | **2.0** | **5.92E-06** |
| *Alcohol metabolism* |  |  |  |  |  |
| *Aldh1a1* | Aldehyde dehydrogenase family 1 subfamily A1 | -0.75 | 0.28 | 0.21 | 0.68 |
| *Cyp2e1* | Cytochrome P450 family 2 subfamily e polypeptide 1 | **-1.54** | **0.003** | -0.74 | 0.14 |

### FGF21, a peptide metabolic hormone, is significantly increased in alcohol-diet-fed L1^LKO^ group compared to the alcohol-diet-fed L1^FLX^ group

Fibroblast growth factor (FGF21) is known to be up-regulated following alcohol-feeding, and has been reported to have a protective role in alcoholic steatohepatitis, both in mice and humans (22, 23). RNA-Seq analyses revealed that *Fgf21* was significantly up-regulated by 4.2-fold (Log2FC=2.0; padj=5.92^E06^) in alcohol-diet-fed L1^LKO^ mice versus the control-diet-fed L1^LKO^ mice, but was unchanged in alcohol-diet-fed L1^FLX^ mice versus the control-diet-fed L1^FLX^ (padj=0.60) (**Table 2**). Consistent with the RNA-Seq data, quantitative PCR also revealed significant upregulation of *Fgf21* mRNA in the alcohol-diet-fed L1^LKO^ group (**Figure 5A**). Next, to test whether secretory levels of FGF21 in the serum were affected in alcohol-diet-fed L1^LKO^ mice, we performed ELISA on serum samples collected from alcohol-diet-fed and control-diet-fed L1^FLX^ and L1^LKO^ mice. The results showed that FGF21 was significantly increased in alcohol-diet-fed L1^LKO^ mice but not in the alcohol-diet-fed L1^FLX^ mice (**Figure 5B**). These data indicated that *Fgf21* may be regulated by ZFP36L1 in this model.

**Figure 5:**
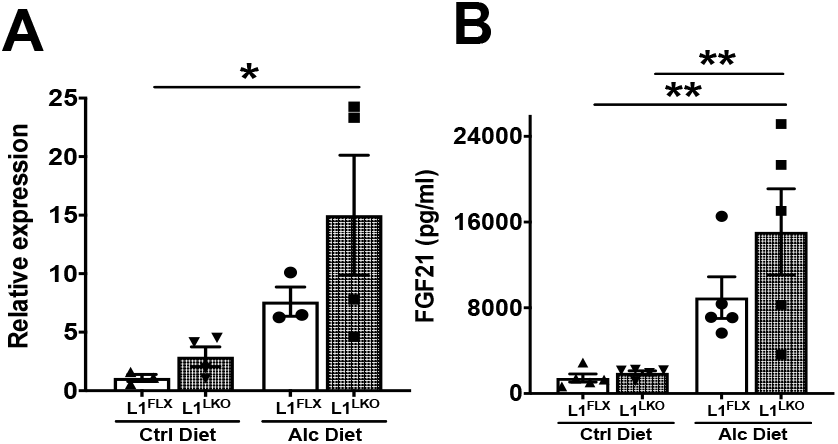
FGF21 is significantly increased in alcohol-diet-fed ZFP36L1 LKO mice. (**A**) Relative expression of *Fgf21* mRNA in the liver homogenate from alcohol-diet-fed ZFP36L1-sufficient (L1^FLX^) and liver-specific ZFP36L1-deficient (L1^LKO^) mice and control-diet-fed ZFP36L1-sufficient (L1^FLX^) and liver-specific ZFP36L1-deficient (L1^LKO^) mice. N=3-4. (**B**) Secreted levels of FGF21 (picograms per milliliter) in the serum of alcohol-diet-fed ZFP36L1-sufficient (L1^FLX^) and liver-specific ZFP36L1-deficient (L1^LKO^) mice and control-diet-fed ZFP36L1-sufficient (L1^FLX^) and liver-specific ZFP36L1-deficient (L1^LKO^) mice. n=5. Statistical analyses were performed by one-way ANOVA followed by Tukey’s correction for multiple comparisons in both (**A**) and (**B**). All data are shown as mean ± SEM. *p<0.05; **p<0.01. FGF21, Fibroblast Growth Factor 21.

### ZFP36L1 regulates *Fgf21* mRNA turnover through AREs in its 3’UTR

Since ZFP36 family members regulate their target mRNAs by binding to their 3’UTR AREs (2), we first tested whether *Fgf21* mRNA contains AREs in its 3’UTR that could be potential ZFP36L1 binding sites. As expected, the 3’UTR of *Fgf21* mRNA contains at least three highly conserved potential ZFP36L1 binding sites (AREs) within its 3’UTR (**Figure 6A).** Next to test whether ZFP36L1 regulates *Fgf21* mRNA, we cloned the complete 3’UTR of *Fgf21* mRNA (total of 163 bp were cloned that also included 37 bp of the protein coding region) into the pMIR-REPORT luciferase reporter construct, and tested whether ZFP36L1 can regulate the stability of *Fgf21* mRNA 3’UTR in luciferase reporter assays. As shown in **Figure 6B**, the un-transfected control (bar 1) or the transfection of beta-galactosidase alone (bar 2) did not result in detectable luciferase activity/luminescence. Similarly, transfection of an empty pcDNA3.1 vector alone (bar 3) or a plasmid expressing wild-type ZFP36L1 (pcDNA3.1-*Zfp36l1*) did not result in detectable luciferase activity. Luciferase activity was reported from cells transfected with the pMIR-REPORT (luciferase reporter expression vector) (bar 5), the pMIR-REPORT containing *Fgf21*-3’UTR (bar 6), a combination of pMIR-REPORT and pcDNA3.1 plasmid expressing WT ZFP36L1 (bar 7), a combination of pMIR-REPORT expressing *Fgf21*-3’UTR and pcDNA 3.1 empty vector (bar 8), and a combination of two empty plasmids (pMIR-REPORT + pcDNA 3.1) (bar 9). Luciferase activity was significantly inhibited (bar 10) when wild-type ZFP36L1 and the pMIR-*Fgf21*-3’UTR were present together, indicating that ZFP36L1 decreased the stability of *Fgf21*-3’UTR. In comparison to this, the luciferase activity was significantly rescued when the ZFP36L1-zinc finger mutant (C135R, C173R) was present (bar 11), indicating that the two intact zinc-fingers are essentially required for this effect. These data show that ZFP36L1 directly regulates *Fgf21*-3’UTR and that this effect is mediated through the two zinc-finger domains of ZFP36L1. **Figure 6C** shows equivalent expression of WT-ZFP36L1 and mutant-ZFP36L1 in HEK293 cells.

**Figure 6:**
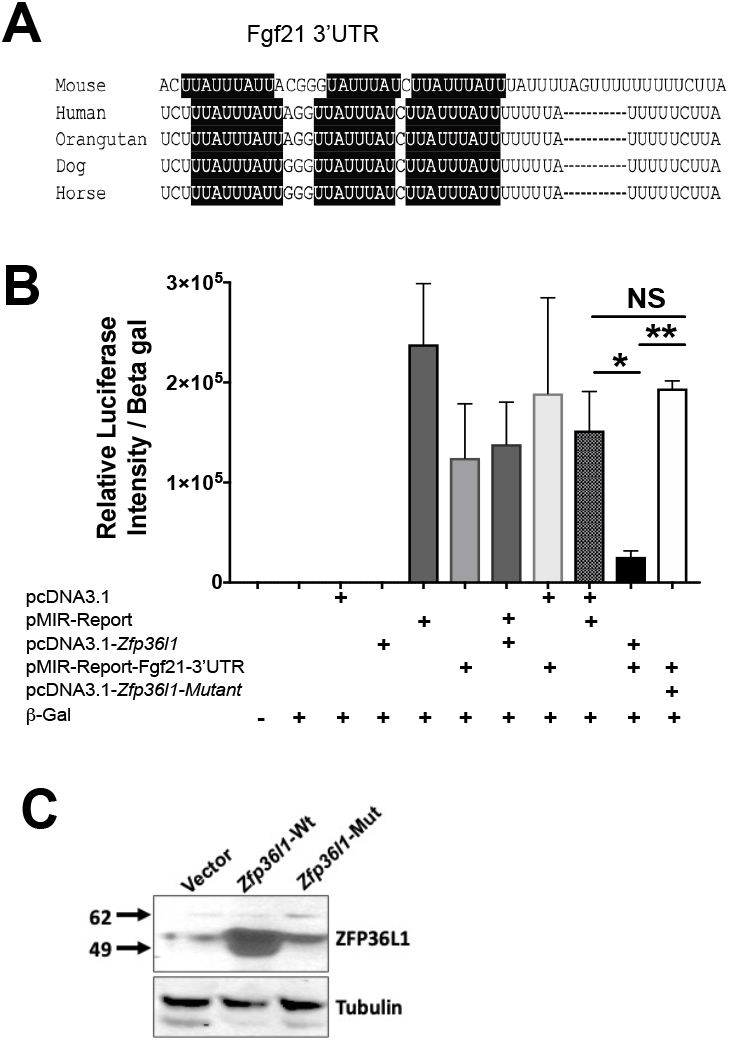
ZFP36L1 regulates the 3’UTR of *Fgf21* mRNA. (**A**) At-least three evolutionarily conserved AU-rich elements (AREs) are present in the 3’-untranslated region of the *Fgf21* transcript. The AREs are highlighted in black. (**B**) HEK293 cells were transfected with empty vector (pcDNA 3.1), luciferase reporter (pMIR-REPORT), WT-human ZFP36L1 expression plasmid (pcDNA 3.1-*Zfp36l1*), ZFP36L1 expression plasmid with C135R, C173R zinc finger mutation (pcDNA 3.1-*Zfp36l1*-mutant), and pMIR-REPORT vector expressing the 3’UTR of *Fgf21* (pMIR-REPORT-*Fgf21-3’UTR*) in various combinations. Beta-galactosidase was transfected as an internal control and data are represented as relative luciferase activity normalized to beta-galactosidase activity. Bar 1: Un-transfected; bar 2: Beta-galactosidase only; bar 3: pcDNA 3.1 empty vector only; bar 4: pcDNA 3.1-*Zfp36l1-wt*; bar 5; pMIR-REPORT empty vector only; bar 6: pMIR-REPORT-*Fgf21*-3’UTR-reporter; bar 7: pMIR-REPORT empty vector + pcDNA 3.1 *Zfp36l1-wt*; bar 8: pcDNA 3.1 empty vector + pMIR-REPORT *Fgf21*-3’UTR-reporter; bar 9: pcDNA 3.1 empty vector + pMIR-REPORT empty vector; bar 10: pcDNA 3.1 *Zfp36l1-wt* + pMIR-REPORT-*Fgf21*-3’UTR; bar 11: pcDNA 3.1 *Zfp36l1*-mutant + pMIR-REPORT *Fgf21*-3’UTR. (**C**) Immunoblot of the cytosolic extracts from cells transfected with vector (pcDNA 3.1) alone (lane 1), pcDNA 3.1 with wild-type human *Zfp36l1* (lane 2) or the zinc-finger mutant (C135R, C173R) form of *Zfp36l1* (lane 3) that were used in the luciferase reporter assay. The position of ZFP36L1 is indicated to the right. The bottom gel shows the loading control, tubulin.

### ZFP36L1 promotes the decay of full-length *Fgf21* mRNA via the zinc-finger motif

We next tested the ability of ZFP36L1 to destabilize a full-length transcript of *Fgf21* through co-transfection assays in HEK293 cells. When a full-length *Fgf21*-mRNA was co-expressed with ZFP36L1-WT protein, the steady state levels of *Fgf21* mRNA were decreased at both 0.25 μg and 0.5 μg concentrations of ZFP36L1 (**Figure 7A, bar 3 and 4**). This effect was statistically significant at 0.5 μg (**Figure 7A, bar 4**). However, when a full-length *Fgf21*-mRNA was co-expressed with zinc-finger ZFP36L1-Mutant (C135R,C173R), the levels of *Fgf21* mRNA did not decrease at either the low (0.25 μg) or the high (0.5 μg) concentration of ZFP36L1-mutant plasmid (**Figure 7A, bar 5 and 6**). In fact, there was an increased accumulation of *Fgf21* mRNA in the presence of ZFP36L1-mutant, particularly at the high concentration (0.5 μg) (**Figure 7A, bar 6**). **Figure 7B** shows the dose-dependent increase in the expression of ZFP36L1-WT and ZFP36L1-mutant in this experiment. This assay suggested that ZFP36L1 promotes the decay of *Fgf21* mRNA and that the zinc-finger motif of ZFP36L1 is required for this effect.

**Figure 7:**
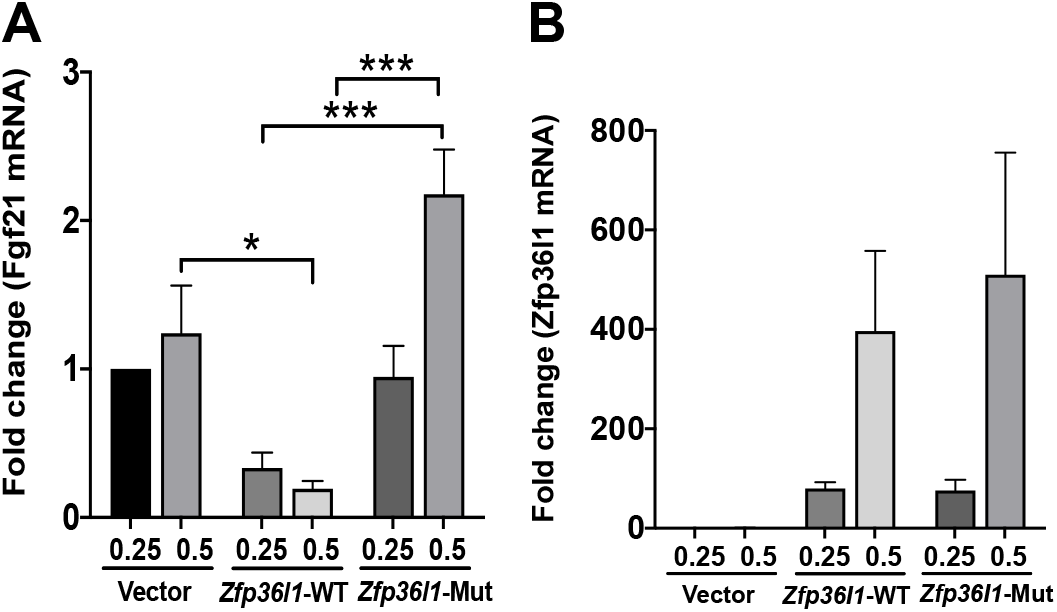
ZFP36L1 reduces the steady state levels of *Fgf21* mRNA. Wild-type *Zfp36l1* or its non-binding mutant (zinc-finger-mutant) plasmids were co-transfected with plasmid expressing full-length *Fgf21* mRNA in HEK293 cells. (**A**) Relative expression of *Fgf21* mRNA 48-hours post-transfection. Bars 1, 2 show *Fgf21* mRNA expression following the transfection of pcDNA 3.1 empty vector at 0.25 and 0.50 μg respectively; bars 3, 4 show *Fgf21* mRNA expression following the transfection of wild-type *Zfp36l1* at 0.25 and 0.50 μg respectively; and bars 5, 6 show *Fgf21* mRNA expression following the transfection of zinc-finger-mutant *Zfp36l1* at 0.25 and 0.50 μg respectively. N=4. (**B**) Relative expression of *Zfp36l1* in HEK293 cells transfected with pcDNA 3.1 empty vector, wild-type *Zfp36l1* expression plasmid, and zinc-finger-mutant *Zfp36l1* expression plasmid. Statistical analyses were done by one-way ANOVA followed by Tukey’s correction for multiple comparisons in both (**A**) and (**B**). All data are shown as mean ± SEM. *p<0.05; ***p<0.001.

### ZFP36L1 binds to *Fgf21*-mRNA 3’UTR AU-rich region

Finally, we tested whether ZFP36L1 expressed in HEK293 cells can directly bind to AREs within the 3’UTR of *Fgf21* mRNA. Biotin 5’-end labelled RNA probes designed from the 3’UTR of *Fgf21* mRNA were synthesized. The probes consisted of a WT *Fgf21*-ARE and a mutant *Fgf21*-ARE in which all the six core A residues were mutated to C residues (**Figure 8A**). Mutations of core A residues to C residues has been previously shown to abrogate the binding of TTP to its target mRNA probes (24). A *Tnf* 3’UTR-ARE probe was used as a positive control. Cell lysates expressing WT ZFP36L1 or zinc-finger mutant ZFP36L1 (C135R, C173R) were incubated with wild-type *Tnf*, wild-type *Fgf21* or mutated-*Fgf21* mRNA ARE probes, and RNA mobility shift assays were performed. As shown in **Figure 8B**, cytosolic extracts from cells expressing vector alone (lane 2) did not bind *Tnf*-ARE. The migration position of the free probe is depicted in **lane 1**. *Tnf*-ARE was able to bind to ZFP36L1, as is evident from the slower migration of the probe (**lane 3**). The *Tnf*-ARE probe was also able to bind to a small amount of zinc finger mutant-ZFP36L1, but with greatly reduced ability (**lane 4**) compared to wild-type ZFP36L1 (**lane 3**), as is evident from the increased migration (downward shifting of the band) in this case (**lane 4**). When the WT *Fgf21*-ARE probe was used, significant binding was evident with the wild-type ZFP36L1 protein (**lane 7**). The bound *Fgf21*-ZFP36L1 complex exhibited significantly reduced migration in the gel, indicating good binding. The binding also appeared to be much better than was observed with the *Tnf*-ARE, as is evident from the larger size and increased intensity of the band (**lane 7**). Again, the zinc-finger mutant-ZFP36L1 was still able to bind the WT *Fgf21*-ARE but with markedly reduced ability (**lane 8**). The position of the wild-type *Fgf21*-ARE free probe is shown in **lane 5**. **Lane 6** shows the vector control. Finally, as shown in **lane 11**, the mutant-*Fgf21*-ARE probe was able to bind to the wild-type ZFP36L1 protein; however, its ability to bind was much less than that of the WT *Fgf21*-ARE probe (**lane 7**). This is clear from increased migration (downward shifting of the band) and reduced intensity and splitting of the band (**lane 11**). The mutant-*Fgf21*-ARE probe was, however, not able to bind to the mutant-ZFP36L1 (**lane 12**). The position of the mutant-*Fgf21*-ARE free probe is shown in **lane 9**. **Lane 10** shows the vector control. **Figure 8C** shows equivalent expression of WT and mutated ZFP36L1 in HEK293 cells.

**Figure 8:**
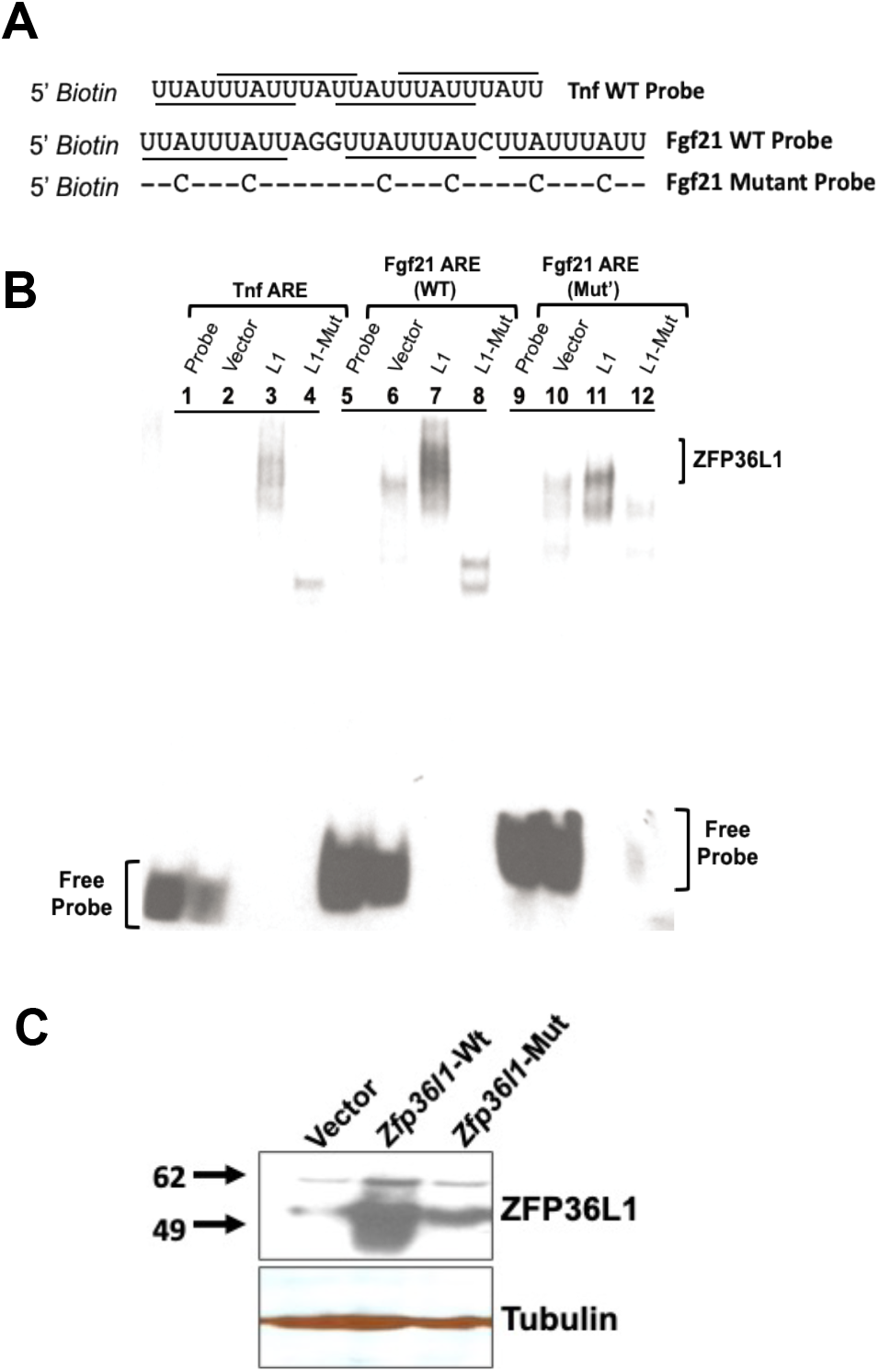
ZFP36L1 binds to the ARE from *Fgf21* mRNA in electrophoretic mobility shift assay. RNA electrophoretic gel-shift mobility assay with ZFP36L1-WT, ZFP36L1-Mutant, WT-*Fgf21*-ARE and Mutant-*Fgf21*-ARE. (**A**) Sequences of 5’-biotin labelled RNA probes corresponding to the WT *Tnf* ARE (GenBank™ accession number X02611), WT *Fgf21* ARE (GenBank™ accession number NM_019113.4), and mutant human *Fgf21* ARE in which six A residues were replaced by C residues). The ARE binding sites (octamer and nonamers) are underlined. The hyphens indicate the sequence identify at that position to the WT-*Fgf21* ARE probe. (**B**) HEK293 cells were transfected with vector alone (pcDNA 3.1), WT-human ZFP36L1 expression construct (pcDNA 3.1-*Zfp36l1*), or the human ZFP36L1 expression construct with C135R, C173R zinc finger mutation (pcDNA 3.1-*Zfp36l1*-mutant). Equal amounts of these cytosolic extracts were combined with a probe derived from *Tnf* ARE (**lanes 1-4**), WT-*Fgf21* ARE (**lanes 5-8**), and Mutant-*Fgf21* ARE (**lanes 9-12**) and gel-shift assay was performed. Lane 1 and 2 shows the position of the free *Tnf* probe when either free probe only (**lane 1**) was used or the probe was combined with empty vector-expressing lysates (**lane 2**). Large complex was formed with the WT-ZFP36L1 (**lane 3**) but not with the mutant-ZFP36L1 (**lane 4**). Lane 5 (probe alone) and 6 (probe with empty vector) shows the position of the free *Fgf21*-WT probe. The amount of probe and the migration position of the complex formed was shifted with WT-ZFP36L1 (**lane 7**) but not with the mutant-ZFP36L1 (**lane 8**). Lane 9 (probe alone) and 10 (probe with empty vector) shows the position of the free *Fgf21*-mutant probe. The amount of probe and the migration positions of the complex formed was partially shifted with WT-ZFP36L1 (**lane 11**) but not with the mutant-ZFP36L1 (**lane 12**). (**C**) Immunoblot of the cytosolic extracts from cells transfected with vector (pcDNA 3.1) alone (**lane 1**), wild-type human ZFP36L1 (**lane 2**) or the zinc-finger mutant (C135R, C173R) form of ZFP36L1 (**lane 3**) that were used in the RNA electrophoretic mobility shift assay. The position of ZFP36L1 is indicated to the right. The bottom gel shows the loading control, tubulin.

## DISCUSSION

Excessive alcohol consumption is a global problem with enormous clinical and economic consequences (25). Liver, the primary site of alcohol metabolism, sustains the greatest tissue injury following alcohol ingestion. However, the pathogenic mechanisms involved in alcoholic hepatic steatosis, injury, and inflammation remain incompletely understood. The current study describes an interesting role of a posttranscriptional regulator of gene expression, i.e., ZFP36L1, in the pathogenesis of alcohol-induced changes in the liver.

Liver-specific ablation of ZFP36L1 (L1^LKO^ mice) resulted in significant reduction in hepatic steatosis, liver injury, and inflammation in response to alcohol when compared to mice that contained an intact *Zfp36l1* gene (L1^FLX^ mice). These findings suggested that ZFP36L1 promoted instability of mRNAs that confer protection against alcohol-induced liver injury. Along similar lines, liver-specific loss of ZFP36L1 was shown to confer protection against diet-induced obesity and steatosis by altering bile acid metabolism through regulation of cholesterol 7α–hydroxylase (Cyp7a1) mRNA (11). Loss of ZFP36L1 was also shown to protect against the development of experimental osteoarthritis through the post-transcriptional regulation of heat shock protein 70 family members (26). Finally, a recent study showed that ZFP36L1 is required for the maintenance of the marginal zone B-cell compartment, and this effect was dependent on its regulation of the transcription factors KLF2 and IRF8 (27). Together, these studies indicate that ZFP36L1 regulates a variety of physiological and pathological responses through its mRNA targets.

Alcoholic hepatitis is a serious consequence of alcoholic liver disease and is accompanied by increased expression of pro-inflammatory mediators that recruit inflammatory cells into the liver. RNA-sequencing revealed that a large number of inflammatory mRNAs were significantly upregulated in L1^FLX^ mice, but not in the L1^LKO^ mice, after alcohol administration. For instance, TNFa levels are increased in both human alcoholic liver and in animals models of liver disease (28). *Tnf* mRNA was significantly upregulated in L1^FLX^ but not in the L1^LKO^ group post-alcohol-challenge. MCP1, also known as CCL2, levels are increased in the plasma of patients with alcoholic liver disease and have been correlated with liver neutrophil infiltrates after alcohol (29). Further, deficiency of MCP1 has been shown to protect against alcoholic liver injury (18). *Mcp1/Ccl2* was significantly upregulated in the L1^FLX^ but not in the L1^LKO^ mice post-alcohol-administration. *Cd14* (a coreceptor for toll-like receptor 4) was specifically upregulated in the alcohol-fed L1^FLX^ group but was unchanged in the alcohol-fed L1^LKO^ group. CD14-deficient mice have been shown to exhibit reduced early alcohol-induced liver injury (30). Liver macrophages and monocytes from alcoholic hepatitis patients show increased activation of NFkB (31). *Ikbip* and *Nfkbib*, two members of NFkB family, were significantly up-regulated in the alcohol-fed L1^FLX^ group but were unchanged in the alcohol-fed L1^LKO^ group. Similarly, *Mapk4*, a member of the MAP kinase signaling family, is upregulated following alcohol exposure, both *in vitro* and *in vivo* (31). While *Mapk4* was significantly up-regulated in the alcohol-fed L1^FLX^ group, its expression levels remained unchanged in the alcohol-fed L1^LKO^ group. Finally, *Mmp12*, a metalloprotease involved in extracellular remodeling (32), was significantly upregulated in the alcohol-fed L1^FLX^ group but not in the alcohol-fed L1^LKO^ group. Of note, *Mmp12* has been shown to be upregulated in primary murine macrophages following alcohol exposure (33). These outcomes suggest that the ZFP36L1-deficient liver cells in alcohol-challenged L1^LKO^ mice exhibit mitigated cellular injury that in turn prevents the pronounced inflammation, a response evident in alcohol-challenged L1^FLX^ mice. Second, we speculate that other members of this protein family, particularly ZFP36/TTP, can regulate inflammation, whereas ZFP36L1 regulates metabolic genes within the liver. These possibilities, however, need to be tested further.

FGF21, a liver-specific metabolic hormone, is known to be elevated in the blood of both mice and humans after acute or binge alcohol ingestion (22, 34). Interestingly, FGF21 was significantly upregulated in the alcohol-fed L1^LKO^ group but not in the alcohol-fed L1^FLX^ group, suggesting that ZFP36L1 directly or indirectly regulates the expression of *Fgf21*. This was also indicated by the finding that PPARa, the transcriptional regulator of FGF21, was significantly downregulated in both the alcohol-fed L1^FLX^ and the L1^LKO^ groups. Accordingly, we conducted further experiments to determine whether *Fgf21* is a direct target of ZFP36L1. First, using luciferase activity assays, we demonstrated that ZFP36L1 can destabilize *Fgf21*-3’UTR. Second, using co-transfection assays, we demonstrated that ZFP36L1 can destabilize the full-length *Fgf21* mRNA. Third, using electrophoretic mobility shift assays, we demonstrated that ZFP36L1 can directly bind to the 3’UTR ARE region of *Fgf21* mRNA. Finally, we established that this effect was mediated by the two tandem zinc-finger-domains of ZFP36L1, as indicated by the reappearance of the luciferase activity and the stabilization of *Fgf21* mRNA when a zinc-finger-mutant of ZFP36L1 replaced the WT-ZFP36L1 in the luciferase experiment and the co-expression assays respectively. These experiments demonstrated that *Fgf21* mRNA can be a target of ZFP36L1.

FGF21 has been previously shown to be a target of ZFP36/TTP, another member of the ZFP36 family of RNA binding proteins. For instance, Sawicki et al. demonstrated that *Fgf21* mRNA is post-transcriptionally regulated by ZFP36/TTP, and that liver-specific loss of ZFP36/TTP improves systemic glucose tolerance and insulin sensitivity in mice fed a high-fat-diet by upregulating FGF21 (35). Another study recently demonstrated that CCR4-NOT deadenylase-ZFP36/TTP complex promotes degradation of *Fgf21* mRNA, and that ablation of CCR4-NOT deadenylase activity decreases the susceptibility of mice to metabolic disorders, including obesity and steatosis, by increasing FGF21 levels (36). Our data provide evidence of redundancy between ZFP36/TTP and ZFP36L1 in the post-transcriptional regulation of *Fgf21* mRNA. However, the relative contribution of ZFP36/TTP versus ZFP36L1 in the posttranscriptional regulation of *Fgf21* mRNA remains unexplored and warrants further investigation.

FGF21 has been shown to confer protection from alcohol-induced hepatic steatosis and liver injury in previous studies. In fact, chronic alcohol feeding in the absence of FGF21 results in significant liver injury in mice (22). Mice lacking FGF21 have been shown to exhibit exacerbated alcohol-induced liver steatosis and liver injury through activation of genes involved in lipogenesis and repression of genes involved in beta oxidation of fatty acids (23). Conversely, administration of recombinant FGF21 to WT mice that were chronically fed alcohol was shown to significantly attenuate hepatic steatosis and liver injury (23). Consistent with these reports, we demonstrate that the upregulation of FGF21 aligns with the hepatoprotective effect in ZFP36L1 liver-specific knockout mice. However, it remains to be tested whether the hepatoprotective effect of ZFP36L1 deletion is mediated through the upregulated expression of FGF21 alone or through the upregulation and downregulation of a battery of anti-inflammatory and proinflammatory genes, respectively.

Previous studies with the ZFP36 RNA binding protein family have suggested that a number of mRNA targets of these proteins are cell-specific. For instance, *Tnf* has been identified as a myeloid-cell-specific target of ZFP36/TTP (37). Similarly, *Notch 1* was identified as a thymocyte-specific target of both ZFP36L1 and ZFP36L2 (38). Our study indicates that *Fgf21* mRNA is a target of ZFP36L1 in hepatocytes/cholangiocytes. However, it remains unclear whether this relationship is specific to one or both the cell types. Further, whether the deletion of ZFP36L1 in other cell types such as immune cells and hepatic stellate cells within the liver will also result in the upregulation of *Fgf21* also warrants further investigation.

Previous studies with ZFP36 RNA binding protein family have also indicated stimulus-specificity for their mRNA targets. For example, myeloid-specific loss of ZFP36/TTP did not result in any abnormalities under basal conditions (39). However, these mice were hypersensitive to lipopolysaccharide challenge that resulted from the stabilization and several-fold up-regulation of ZFP36/TTP target, *Tnf* (39). Similarly, TTP deletion in keratinocytes resulted in hypersensitivity to imiquimod-induced skin inflammation in these mice and this effect was mediated by keratinocyte-specific stabilization and upregulation of *Tnf* (40). In a recent study, genetic ablation or silencing of ZFP36L1 significantly protected mice from experimental-osteoarthritis by stabilizing members of the heat shock protein 70 family, again indicating stimulus-specific targets and resulting pathological outcomes (26). Along these lines, *Fgf21* also appears to be a stimulus-specific physiologically relevant target of ZFP36L1.

Collectively, this study reveals a novel post-transcriptional mechanism of regulation of FGF21 production during alcohol-induced hepatic disease. First, ZFP36L1 deletion in the liver resulted in significant reductions in hepatic steatosis, liver injury, and inflammation in response to alcohol. Second, using biochemical approaches, we demonstrated that ZFP36L1 regulates the expression of FGF21, following basic mechanism through which TTP family members operate to regulate their target mRNAs. Of note, FGF21 is considered as a potential treatment of metabolic disease; therefore, various FGF21 analogs have been developed and are undergoing clinical trials for metabolic disorders, including diabetes and obesity (41). Our study suggests that the hepatic ZFP36L1-FGF21 axis is a major signaling route in alcohol-induced hepatic steatosis in mice. This opens up the possibility of silencing ZFP36L1 as a possible method of protection against alcoholic-liver disease.

## Acknowledgments

We thank Sherry Ring for histological tissue processing.

## Author Contributions

S.P. conceived and designed the study; S.P. and C.B. maintained the animal colony. C.B genotyped the animals and performed all the experiments unless mentioned otherwise. C.B. and J.C performed the luciferase assays. J.C. performed the co-transfection and the EMSA assays. Y.S performed the oral gavages. P.J.B provided the *Zfp36l1* floxed mice and edited the manuscript; S.P. performed the histopathological studies; S.P and Y.S. wrote and reviewed the manuscript for intellectual contents.

---

**Supplemental Figure 1:**
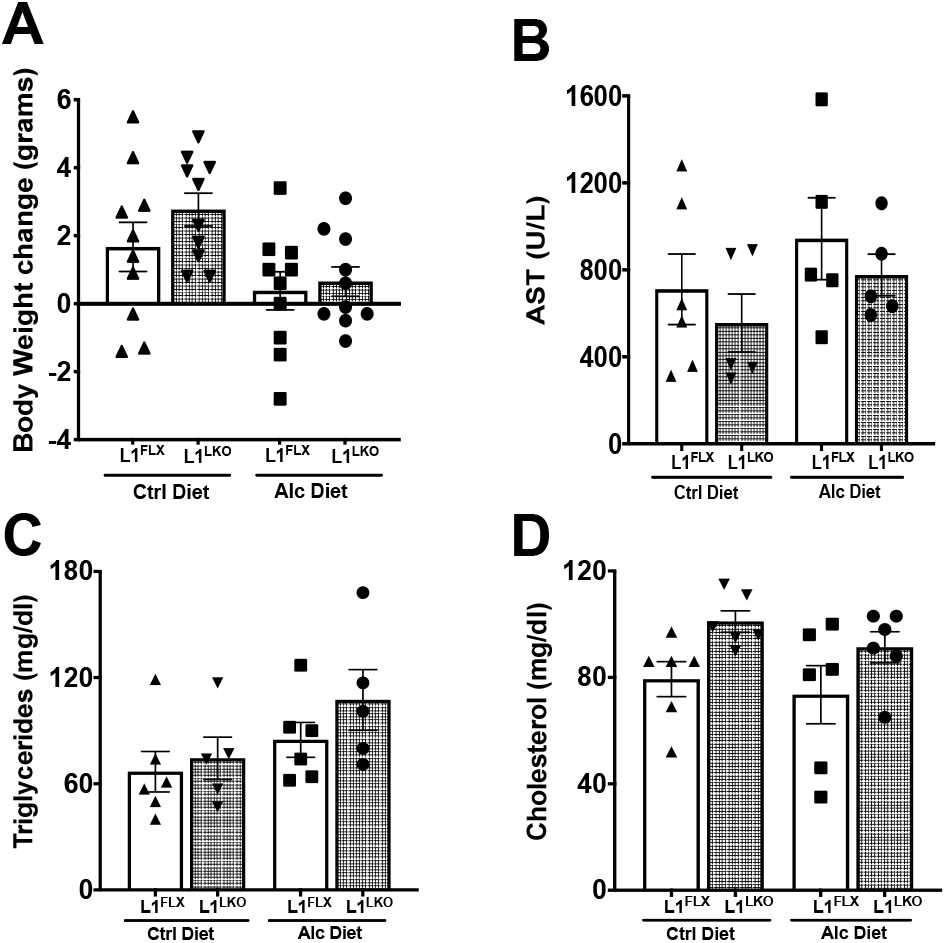
**A.** Change in body weight (initial weight on day 0 was subtracted from final weight on day 11) in both ZFP36L1-sufficient (L1^FLX^) and liver-specific ZFP36L1-deficient (L1^LKO^) mice maintained on alcohol-diet or control-diet. N=10 in each group. **B, C, and D.** Levels of AST, triglycerides, and cholesterol in the serum of alcohol-diet-fed ZFP36L1-sufficient (L1^FLX^) and liver-specific ZFP36L1-deficient (L1^LKO^) mice and the control-diet-fed ZFP36L1-sufficient (L1^FLX^) and liver-specific ZFP36L1-deficient (L1^LKO^). n=5 (ethanol-diet group); n=6 (control diet group).

